# SMuRF: a novel tool to identify genomic regions enriched for somatic point mutations

**DOI:** 10.1101/271957

**Authors:** Paul Guilhamon, Mathieu Lupien

## Abstract

**Motivation:** Single Nucleotide Variants (SNVs), including somatic point mutations and Single Nucleotide Polymorphisms (SNPs), in noncoding cis-regulatory elements (CREs) can affect gene regulation and lead to disease development (Zhou *et al.*, 2016; Zhang *et al.*, 2014). Others have previously developed methods to identify important clusters of somatic point mutations based on proximity (Weinhold *et al.*, 2014) or the enrichment of inherited risk-SNPs at CREs (Ahmed *et al.*, 2017). Here, we present SMuRF (Significantly Mutated Region Finder), a user-friendly command-line tool to identify these significantly mutated regions from user-defined genomic intervals and SNVs.

**Results:** SMuRF identified 72 significantly mutated CREs in liver cancer, including known mutated gene promoters as well as previously unreported regions.

**Availability:** The source code for SMuRF is open-source and freely available on GitHub (https://github.com/LupienLabOrganization/SMuRF) under the GNU GPLv3 license. SMuRF is implemented in Bash and R; it runs on any platform with Bash (≥4.1.2), R (≥3.3.0) and BEDTools (≥2.26.0). It requires the following R packages: GenomicRanges, gtools, gplots, ggplot2, data.table, psych, and dplyr.

**Supplementary Information:** Supplementary information available at Bioinformatics online.

## 1 INTRODUCTION

With the advent of next-generation sequencing technologies, a growing catalogue of genome-wide datasets has become available. This includes whole-genome sequencing to detect single nucleotide variants (SNVs) in diseased tissue (eg: TCGA Research Network: http://cancergenome.nih.gov/) as well as maps of histone variants and chromatin accessibility (ENCODE Project Consortium, 2012). Using these datasets, numerous CREs have been identified as recurrently mutated in cancer and other diseases. A notable example is the *TERT* promoter in glioma, melanoma, medulloblastoma, hepatocellular carcinoma, lung adenocarcinoma, thyroid and bladder cancers (Vinagre et al., 2013). The mutations in this promoter create new transcription factor binding sites, leading to increased *TERT* expression and lengthening of telomeres contributing to oncogenesis. Enhancers and anchors of chromatin interaction can also display recurrent mutation, such as the *PAX5* enhancer in chronic lymphocytic leukemia (Cobaleda et al., 2007; Puente et al., 2015) and CTCF binding sites in colorectal cancer (Katainen et al., 2015).

As the function of a CRE can be disrupted by SNVs at numerous positions, recurrently mutated CREs must be identified by comparing the rate of mutation of a given element to that of all other regulatory elements in the same cell or tissue type. Weinhold *et al* proposed such a method to identify noncoding regions enriched for somatic point mutations (Weinhold et al., 2014). It relied on an initial clustering of nearby SNVs to determine regions of interest. A binomial test was then used to determine whether a given region had a higher burden of mutation than could be expected based on the mutation rate of the other clusters of SNVs. Here we have expanded this methodology with our Significantly Mutated Region Finder (SMuRF) by using a user-defined set of regions of interest as the input rather than relying on a proximity clustering of SNVs and providing a user-friendly tool to identify, filter, and annotate significantly mutated genomic regions.

## 2 METHODS

SMuRF consists of two main steps. The first filters, counts, annotates, and intersects the list of SNVs with the set of genomic coordinates, using a custom Bash script and the BEDTools suite (Quinlan and Hall, 2010). The second consists in running a binomial test in R followed by a mutation rate filter to determine which genomic intervals are significantly enriched in SNVs and producing output figures as well as files for downstream analyses.

### 2.1 Input processing

The SNVs in BED or vcf format, are optionally filtered for known SNPs. This will remove either all known SNPs or only those with a minor allele frequency above 1% to preserve potentially interesting acquired SNVs that also occur as extremely rare polymorphisms in the population.

Subsequently, the input genomic regions are annotated as either gene promoter regions or as distal regulatory elements. This is done by overlapping those genomic intervals with a catalogue of gene promoters, derived from Gencode transcription start site annotations (Harrow et al., 2012).

Finally, the input SNVs and genomic intervals are intersected to map all SNVs to unique genomic intervals, and the resulting data structure forms the starting point of the statistical analysis for mutation enrichment.

All of the above filtering and annotating can be achieved with data from any genome for which the required annotation files are available. Those for human builds *hg19* and *hg38* are supplied with the tool for convenience.

### 2.2 Identifying significantly mutated regions

The binomial test used by SMuRF to determine whether a given genomic region is significantly enriched for mutations requires an expected mutation rate. Depending on the sample cohort, the user can choose how this mutation rate is calculated. For each sample, the average number of mutations per base pair in input regions is calculated first. The “*allsamples*” option uses the average of those individual mutation rates across the entire sample cohort. However, if a subset of samples is more or less mutated than the rest, this could lead to biased results when a particular region contains mutations from that subset. In these cases, the “*regionsamples*” option can be used, and the expected mutation rate when testing a particular region will be the average of the mutation rates for the individual samples mutated within that region only.

In both cases, the resulting p-value is then adjusted for multiple testing and the final set of regions is further filtered to include only those that pass a mutation rate threshold. This threshold is defined for each cohort by ranking the mutation rates for each region and identifying the inflection point, as previously described <u>(Whyte et al. 2013)</u>.

A number of output files are generated and these are detailed within the manual; they include a list of genes whose promoters are significantly mutated for use in gene ontology analyses, as well as a bed-formatted list of mutated regions annotated as distal regulatory elements to allow the user to associate them to target genes through GREAT (McLean et al., 2010) or C3D (Mehdi et al., 2017). The main output figure is a scatter plot of-log10(q-value) against the number of unique samples mutated in the region, and color-coded to distinguish gene promoters from distal regulatory elements.

## 3. RESULTS

To illustrate the above steps, we used publicly available acquired SNVs from 88 liver cancer samples (Alexandrov et al., 2013) and chromatin accessibility data from HepG2 (ENCODE Project Consortium, 2012) that provides a reference set for CREs. The total number of SNVs per sample used in the analysis after filtering ranged from 1,344 to 25,121 (Supplementary Figure 1A), with an average of 1.2% falling within one of the 278,135 CREs (Supplementary Figure 1B) as identified in HepG2. While the input SNV numbers covered a wide range, no subset of patients was abnormally hyper or hypomutated, so we selected the “*allsamples*” mode to calculate the background mutation rate for each CRE. In total, 9,485 individual CREs contained at least one mutation, of which 72 (6 promoters and 66 distal regulatory elements) were found to be significantly enriched for mutations (q-value ≤ 0.05 and peak mutation rate ≥ threshold) (Figure 1 and Supplementary Table 1). These regulatory elements were each recurrently mutated in 2-5 samples.

**Figure 1:**
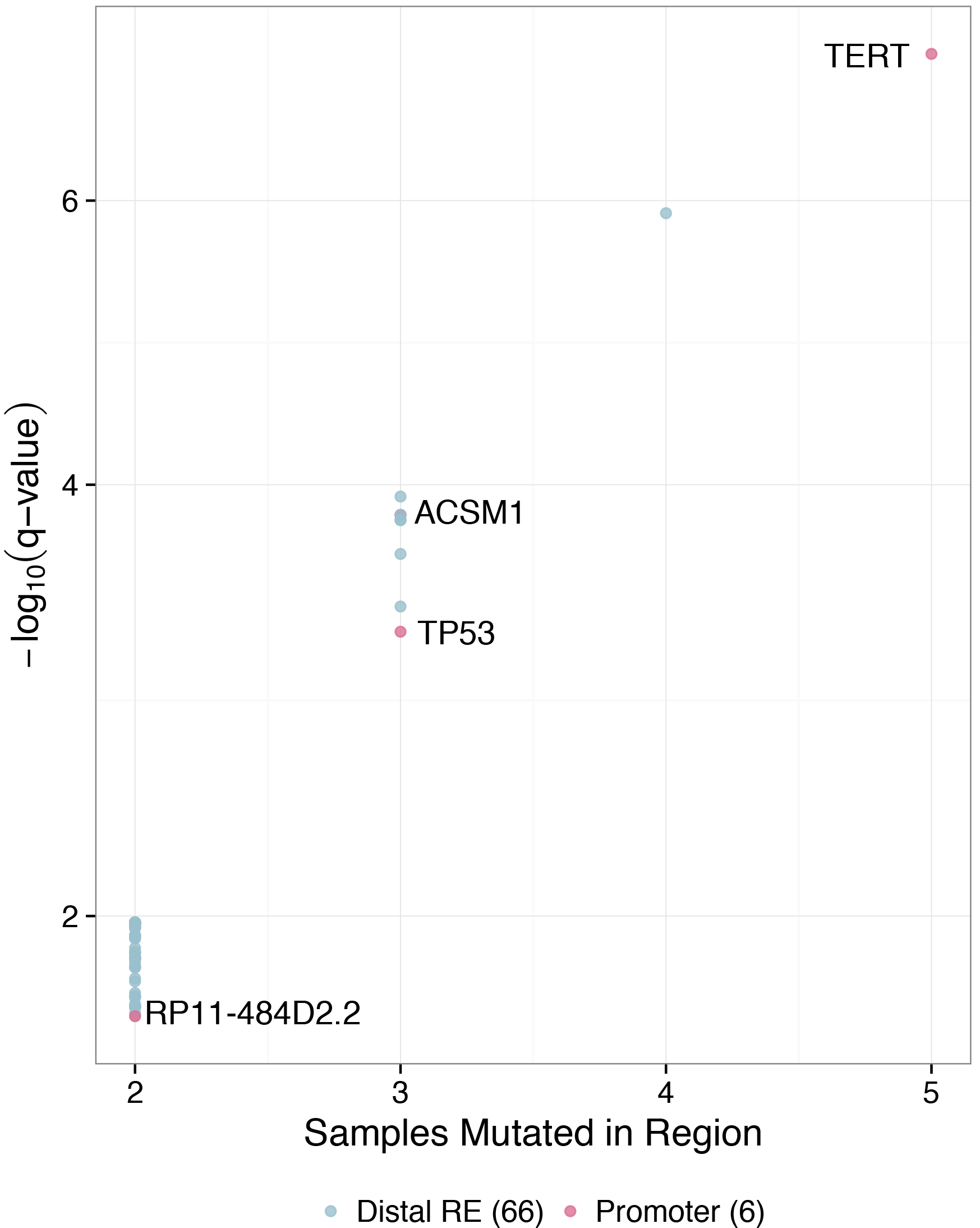
Significantly mutated regions identified by SMuRF. Each of the 72 genomic intervals that passed the significance (q-value ≤ 0.05) and mutation rate filters are represented. The negative log of the q-value calculated from the binomial test for each region is plotted against the number of unique samples with a mutation within that region. The most frequently and most significantly mutated regions include the promoters of both known and novel genes of interest in liver cancer.

Among the highly mutated promoters were those for the *TERT*, *TP53*, *ACSM1*, *TNFRSF8*, and *PCGF5* genes, all previously reported recurrently mutated regions in liver cancer (Fujimoto et al., 2016). Also significantly mutated, however, was the promoter of a gene with unknown function, *RP11-484D2.2*, highlighting the potential of this type of analysis for uncovering novel regions of interest.

Whole-genome sequencing and chromatin accessibility data sets in numerous normal and diseased tissues are becoming more commonly available. SMuRF aims to help further our understanding of the importance of non-coding elements in disease initiation and progression, by highlighting those regulatory elements most likely to have a functional importance due to their high burden of mutation.

## ACKNOWLEDGEMENTS

*Conflict of Interest*: none declared.

